# The ghr-miR164 and *GhNAC100* module participates in cotton plant defence against *Verticillium dahliae*

**DOI:** 10.1101/440826

**Authors:** Guang Hu, Yu Lei, Le Wang, Jianfen Liu, Ye Tang, Zhennan Zhang, Aiming Chen, Qingzhong Peng, Zuoren Yang, Jiahe Wu

**Author notes:** G. H., Y. L., L. W., contributed equally to this study. Corresponding author: J. W., Z. Y. **Authors emails:**Guang Hu, Yu Lei, Le Wang, Jianfen Liu, Ye Tang, Zhennan Zhang, Aiming Chen, Qingzhong Peng, Zuoren Yang, Jiahe Wu.

## Abstract

Previous reports have shown that many miRNAs were identified at the early induction stage during which *Verticillium dahliae* localizes at the root surface. In this study, we constructed two sRNA libraries of cotton root responses to this fungus at the later induction stage when the pathogen enters the root vascular tissue. We identified 71 known miRNAs and 378 novel miRNAs from two pathogen-induced sRNAs and the control libraries. Combined with degradome and sRNA sequencing, 178 corresponding miRNA target genes were identified, in which 40 target genes from differentially expressed miRNAs were primarily associated with oxidation-reduction and stress responses. More importantly, we characterized the ghr-miR164-GhNAC100 module in the response of the plant to *V dahliae* infection. A GUS fusion reporter showed that ghr-miR164 directly cleaved the mRNA of *GhNAC100* in the post-transcriptional process. ghr-miR164-silencing increased the resistance of the plant to this fungus, while the knockdown of *GhNAC100* elevated the susceptibility of the plant, indicating that ghr-miR164-GhNAC100 modulates plant defence through the post-transcriptional regulation. Our data documented that there are numerous miRNAs at the later induction stage that participate in the plant response to *V. dahliae*, suggesting that miRNAs play important roles in plant resistance to vascular disease.

**Highlight:** According to degradome and sRNA sequencings of cotton root in responses to *Verticillium dahliae* at the later induction stage, many miRNAs and corresponding targets including ghr-miR164-GhNAC100 module participate plant defence.

## Introduction

Cotton (*Gossypium hirsutum* L.) is a vital textile and oil crop in the world, but its productivity is constrained by various biotic and abiotic stresses (Xie *et al.*, 2015). One of the stresses is Verticillium wilt, which is a highly destructive vascular disease primarily caused by the soil-borne fungus *Verticillium dahliae* Kleb (Bhat and Subbarao, 1999). The representative symptoms of diseased cotton plants include leaf curl, necrosis and defoliation, and stem wilt (Sink and Grey, 1999). *V. dahliae* generally enters into the vascular tissue through wounded root sites and colonizes and grows in xylem vessels and other dead cell tissues (Bejaranoalcazar *et al.*, 1997; Klosterman *et al.*, 2009). Although no disease symptoms are evident at the stage of pathogen colonization in the xylem vessels, molecular mechanisms, including physiological and biochemical status, should result in remarkable changes in the root cells, especially those around the vascular tissues, resulting from a substantial amount of gene expression reprogramming at the transcription and translation levels.

microRNA (miRNA) is an important component in the post-transcriptional regulation of the target gene expression, playing major roles in plant development and stress responses (Jones-Rhoades *et al.*, 2006). miRNAs can recognize corresponding mRNA targets based on sequence complementarities and guide the direct cleavage of target mRNAs and/or translational repression (Li *et al.*, 2013). Recently, miRNA-mediated gene silencing was found to play a significant role in plant defence against pathogens (Khraiwesh *et al.*, 2012; Shriram *et al.*, 2016). For example, Arabidopsis miR393 was the first miRNA discovered to be involved in plant immunity (Navarro *et al*, 2006). Overexpressing both miR160a and miR398b in rice *(Oryza sativa)* increased the resistance of the plant to *Magnaporthe oryzae* as demonstrated by decreased fungal growth and the upregulated expression of defence-related genes in transgenic rice plants (Li *et al.*, 2014). In cotton plants infected with the fungus, the production of miR166 and miR159 was increased and outputted into the fungal hyphae of *V. dahliae* for specific silencing (Zhang *et al.*, 2016). When miR482 was silenced in the cotton plants, the expression of the *NBS-LRR* defence genes was upregulated, resulting in increasing resistance to fungal pathogen attack (Zhu *et al.*, 2013). Wang *et al.* (2017a) reported that the ghr-miR5272a-mediated regulation of *GhMKK6* transcription contributes to the cotton plant immune response. The evidence is demonstrated in the participation of the cotton miRNAs in plant defence, but the molecular mechanisms and mode of regulation of miRNAs and their corresponding target genes are still unclear.

miRNAs directly participate in various classes of gene expression by post-transcriptional regulation. Among those genes, many transcriptional factors, including NAC, MYB and WRKY, are post-transcriptionally regulated by miRNA, repressing/promoting the expression of downstream genes (Schwechheimer *et al.*, 1998; Hao *et al.*, 2012; Yu *et al.*, 2012). The name of NAC comes from acronym of NAM (no apical meristem), ATAF (Arabidopsis transcription activation factor) and CUC (cup-shaped cotyledon), which contains a miR164 complementary site with few mismatches (Ooka *et al*., 2003; Nuruzzaman *et al*., 2010). NAC is negatively regulated by miR164 to participate in development and defence (Baker *et al.*, 2005; Sieber *et al.*, 2007). In Arabidopsis, miR164 targets the transcripts of six *NAC* genes and prevents organ boundary enlargement and the formation of extra petals during flower development (Laufs *et al*., 2004; Mallory *et al*., 2004). ORE1, an NAC transcription factor, is involved in leaf cell death through miRNA164 regulation (Kim *et al.*, 2009). The miR164 function in the responses of plants to biotic stresses has been verified through the regulation of its corresponding target genes (Bazzini *et al.*, 2007; Bazzini *et al*., 2009; Jia *et al.*, 2009; Xin *et al.*, 2010; Zhao *et al.*, 2012; Feng *et al.*, 2014). Among these target genes, TaNAC21/22 participated in the resistance of wheat plants to stripe rust regulated by tae-miR164 (Feng *et al.*, 2014).

miRNA expression sequencing in the early response of the cotton plants to *V. dahliae* infection has been reported, which showed that many early induction miRNAs were observed that possibly participated in plant defence (Yin *et al.*, 2012; He *et al*., 2014; Zhang *et al.*, 2015a). For example, Yin *et al.* (2012) investigated the transcriptional profile of the miRNAs in Verticillium-inoculated cotton roots at 12 and 24 hours and identified 215 miRNA families and 14 novel miRNAs. Two small RNA (sRNA) libraries were constructed from the seedlings of the upland cotton variety KV-1 inoculated with *V. dahliae* at 24 and 48 hours; 37 novel miRNAs were identified, and potential target genes of these miRNAs were predicted (He *et al.*, 2014). Zhang *et al.* (2015a) conducted sRNA sequencing and degradome sequencing of cotton roots inoculated with *V. dahliae* at 24 hours and identified 140 known miRNAs and 58 novel miRNAs. However, in the later stage of fungal-infected plants (*V. dahliae* has colonized in xylem vessels), the response of miRNA expression has not been investigated using miRNA sequencing.

In this study, we investigated the sRNA expression profiles in cotton roots inoculated by *V. dahliae* at 7 and 10 days and analysed the difference in the expression of known and novel miRNAs compared to mock-treated plants, as well as the functional analysis of the corresponding target genes through sRNA high-throughput and degradome sequencing. The results showed that 71 known miRNAs and 378 novel miRNAs were identified from three sRNA libraries, and 40 corresponding target genes from differentially expressed miRNAs were found by coupling with degradome sequencing. In the late stage of the roots infected by *V. dahliae*, many miRNAs showed a significant difference in their expression level compared to the mock-treated roots. Among these differentially expressed miRNAs, gh-miR164 and its target gene *GhNAC100* were found to form a module to participate in the plant resistance to *V. dahliae* through genetic and biochemical analyses. These findings reveal a miRNA-mediated regulatory network with a critical role in the plant response of pathogen infestation in the main battlefield of vascular tissues.

## Materials and methods

### Plant growth condition and treatment

*G. hirsutum* cv. Jihe713 was donated by Prof. Xiaoli Luo from the Institute of Cotton Research, Shanxi Academy of Agricultural Science. Seedlings grew in a greenhouse at 28°C under a 16-h light/8-h dark photoperiod.

To conduct high-throughput sRNA and degradome sequencing at the two-leaf stage, cotton plants under hydroponic conditions were inoculated with *Verticillium dahliae* strain V991. The 7- and 10-day post inoculation roots, as well as the mock-treated control, were harvested. The inoculated roots were treated ultrasonically (30 seconds with a gap of 30 seconds, repeated 5 times) to remove the fungal hyphae and conidia on their surface. Three types of samples were immediately frozen in liquid nitrogen and stored at −80°C prior to the RNA isolation. The same experiment was repeated twice.

*Nicotiana benthamiana* plants were grown in the greenhouse under a 16-h light/8-h dark photoperiod at 23°C for gene transient expression analyses.

### Fungal cultivation and inoculation

*V. dahliae* strain V991, a strongly pathogenic defoliating isolate, was cultured on potato dextrose agar (PDA) media for a week at 25°C. The mycelia were transferred into Czapek-Dox media for a week at 25°C with shaking (180 rpm) to collect the conidia. For *V. dahliae* infection, the roots of cotton plants were dipped with a conidial suspension (10^6^ conidia mL^−1^) for 50 min. Subsequently, the plants were transferred into fresh, steam-sterilized water for culture or planted into the pot with soil.

### RNA extraction and qPCR analysis

Total RNA was isolated from cotton samples using the PureLink Plant RNA Reagent (Life Technologies, USA) according to the manufacturer’s instructions. First-strand cDNA was synthesized using an EasyScript First-Strand cDNA Synthesis SuperMix (TransGen, Beijing, China). miRNA first-strand cDNA synthesis, qPCR analysis and primer design were conducted as described by Varkonyi-Gasic *et al.* (2007). The qPCR experiment was performed using a TransStart Top Green qPCR SuperMix Kit (TransGen, Beijing, China) in a 20 μL reaction volume on a CFX96TM Real-time Detection System (Bio-Rad Laboratories, Inc., Hercules, Calif). The PCR programme was as follows: pre-denaturation at 95 °C for 30 s, 40 cycles of 95 °C for 15 s, 55 °C for 15 s and 72 °C for 15 s, and a melt cycle from 65 to 95 °C. The 2 ^−ΔΔCT^ method was used to determine the relative expression levels of the miRNAs and target genes. The *UBQ7*gene from *G. hirsutum* was used as an internal control.

Fungal biomass quantification with qPCR techniques was performed as described previously (Wang *et al.*, 2017b). The primer pairs to detect the *V. dahliae* β-*tubulin* gene and the cotton gene *Actin* were used for qPCR. The same experiment was conducted using three biological replicates. The primers used for qPCR are listed in Supplementary Table S11.

### Construction of the sRNA and degradome libraries

The cotton sRNA libraries were constructed using an NEB Next, Ultra sRNA Sample Library Prep Kit (Illumina, San Diego, CA, USA) according to the manufacturer’s instructions. sRNA was purified from 1.5 μg of the total RNA by using the sRNA Sample Pre Kit and ligated first to a 5’ RNA adaptor and then to a 3’ RNA adaptor using T4 RNA Ligase 1 and T4 RNA Ligase 2 (truncated). Reverse transcription synthesis cDNA was purified by polyacrylamide gel electrophoresis as the sRNA library. The sRNA libraries were subjected to high-throughput sequencing with HiSeq2500 (Illumina, San Diego, CA, USA) with a read length of single-end (SE) 50 nt at the Biomarker Technologies Company in Beijing.

The cotton degradome libraries were constructed as previously described (German *et al*., 2008). Briefly, a 5’ RNA adaptor with a *Mme* I recognition site at the 3’ end was ligated to the resulting 42 bp (base pair) fragments consisting of a free phosphate at the 5’ end followed by reverse transcription to cDNA. After PCR amplification, they were digested by the enzyme *Mme* I and ligated to an Illumina 3’ TruSeq adaptor, followed by PCR amplification with a library-specific index primer and a common 5’ primer for multiplex sequencing, gel-purified, and subjected to sequencing by synthesis (SBS) using HiSeq2500 (Illumina, USA) at the Biomarker Technologies Company in Beijing.

### Identification and analysis of known and novel miRNAs

Raw sequences obtained from the three sRNA libraries were first cleaned by filtering out low-quality tags, poly(A) tags, and tags with 3’ adaptor nulls, insert nulls, 5’ adaptor contaminants, or those smaller than 18 nt. Using Bowtie tools soft (Langmead *et al.*, 2009), ribosomal RNA (rRNA), transfer RNA (tRNA), small nucleolar RNA (snoRNA) and other ncRNA and repeats were filtered from the clean reads, respectively, with the Silva, GtRNAdb, Rfam and Repbase database sequence alignments. The remaining sequences from 18~30 nt long were used for miRDeep2 and miRBase (http://www.mirbase.org/ftp.shtml) to identify conserved miRNAs and novel 5p- and 3- derived miRNAs (Friedlander *et al.*, 2012; Zhang *et al.*, 2015b). Only the sequences that were ≤ 2 mismatches with known miRNAs were considered as conserved miRNAs. Otherwise, the reads were defined as non-conserved reads. Unannotated reads were used to predict novel miRNAs based on the characteristic hairpin structure of the microRNA precursors using miRDeep2 (Friedlander *et al.*, 2012). The miRDeep2 software was used to sequence the unannotated reads with the reference genome (TM-1 v1.1, http://mascotton.njau.edu.cn/html) to obtain the positional information on the reference cotton genome, which are mapped reads.

### Differential expression analysis

The reads of each library were normalized by TPM (Transcript per million), normalized expression = (actual miRNA count/total count of clean reads) × 1,000,000 (Fahlgren *et al*., 2007). Differential expression analysis of the inoculated root libraries compared to the control was performed using the DESeq R package (Anders and Huber, 2010). To investigate differentially expressed miRNAs between the treated libraries and the control, the fold change of each identified miRNA was calculated as the ratio of read counts in the treatment libraries to the read counts in the control library followed by the transformation of log2. The value of the log2 Ratio ≥1 or ≤−1, indicating the ratio of fold change (FC) values for the treatments and control libraries, were considered to be significantly differentially expressed. To show the differential expression profiles, heatmaps and clusters were constructed for the miRNAs using ImageGP (http://www.ehbio.com/ImageGP/index.php/Home/Index/index.html).

### Identification of the miRNAs targets by degradome sequencing

The sequences of clean full-length reads collated from the degradome sequencing were used for subsequent analysis after removing low quality sequences and adaptors. There were no mismatches allowed on the 10^th^ and 11^th^ nucleotides of the mature miRNAs where the splice site on miRNA targets generally occurs during degradome analysis. A potential miRNA target with a *P*-value of <0.05 by PAREsnip software was retained, and T-plot figures were drawn. All the target sequences were categorized into five classes based on the abundance of the degradome tags indicating miRNA-mediated cleavage. Category 0-4 was determined as previously described (Liu *et al*., 2014).

### Function enrichment analysis

The miRNA targets in the plants were predicted with TargetFinder software (Allen *et al.*, 2005). Gene Ontology (GO) enrichment analysis of the target genes corresponding to the miRNAs and differentially expressed miRNAs was implemented with GOseq R packages based on Wallenius non-central hyper-geometric distribution (Ashburner *et al*., 2000).

KEGG (Kyoto Encyclopedia of Genes and Genomes, Kanehisa *et al.*, 2004) is a database resource to understand high-level functions and utilities of the biological system, such as the cell, the organism and the ecosystem, from molecular-level information, especially large-scale molecular datasets generated by genome sequencing and other high-throughput experimental technologies (http://www.genome.jp/kegg/). We used KOBAS software (Mao *et al.*, 2005) to test the statistical enrichment of differential expression genes in KEGG pathways.

### Phylogenetic analysis

The NAC genes in this study were retrieved from the NCBI data and aligned with the Clustal X programme. Neighbour-joining (NJ) phylogenetic trees were constructed in MEGA 5.2 with 1,000 bootstrap replicas (Tamura *et al*., 2011).

### Gene isolation and vector construction

To elucidate the miR164 post-transcriptional regulating *GhNAC100* expression, *GhNAC100* was isolated from *G. hirsutum* cv. Jihe713, and the target sequence of *GhNAC100* was mutated by PCR methods and designated *GhNAC100T*^mu^. *GhNAC100* and *GhNAC100^mu^* were inserted into pBI121, respectively, resulting in the pBI121-GhNAC100:GUS and pBI121-GhNAC100^mu^:GUS vectors. Cotton *MIR164*, the ghr-miR164 precursor sequences, was isolated and inserted into the pBI121 instead of the *gus* gene, constructed in pBI121-pre-miR164.

For the virus-induced gene silencing (VIGS) analysis, *tobacco rattle virus* (*TRV*)-based vectors, including pTRV1 (pYL192), pTRV2 (pYL156) and pTRV2e were used in this study. TRV:PDS was employed as a positive control vector in the silenced plants, which had been previously reported by Pang *et al.* (2013). The construction of the TRV-related vectors was performed as described by Liu *et al.* (2004) and Sha *et al.* (2014). Briefly, a small tandem target mimic (STTM) sequence of ghr-miR164 containing two imperfect ghr-miR164 binding sites separated by a 48-bp spacer with the restriction enzyme sites Kpn I and Xma I at the 5’ and 3’ ends, respectively, was designed and inserted into the pTRV2e vector to generate the TRV:STTM164 vector (Supplementary Table S11). A *GhNAC100* fragment was isolated and inserted into pTRV2, and the resulting vector was designated TRV:GhNAC100. All the plasmids were transformed into *A. tumefaciens* strain GV3101 using electroporation. All the primers associated with vector construction are listed in Supplementary Table S11.

### Gene transient expression analysis in *N. benthamiana* leaves

Agrobacterium cells grown overnight at 28°C in lysogeny broth (LB) media containing 50 μg mL^−1^ kanamycin, 50 μg mL^−1^ gentamicin and 50 μg mL^−1^ rifampicin were collected and resuspended in infiltration media (MMA buffer, 10 mM MgCl_2_, 10 mM MES-NaOH, and 200 μM acetosyringone; OD600=0.8). After 3 h of incubation, the suspensions were infiltrated into the *N. benthamiana* leaves using a 2 mL needleless syringe.

### GUS activity analysis

At 48 hours after agro-infiltration, the treated leaves were detached, and β-Glucuronidase (GUS) staining analysis was performed as described by Jefferson *et al.* (1987). GUS activity was quantified by 4-methylumbelliferone (4-MU) testing methods as described by Jefferson *et al.* (1987).

### Analysis of VIGS

Agrobacterium culture and treatments were the same as the method of Agrobacterium co-transformation in tobacco described above. Agrobacterium cells containing *TRV:GhNAC100* or *TRV:STTM164* were mixed with an equal amount of Agrobacterium cells with pTRV1 (pYL192). The mixed Agrobacterium cells were agro-inoculated into the fully expanded cotyledons of the cotton seedlings using a sterile needleless syringe. After 12 hours incubation in darkness, the cotton seedlings were transferred to the greenhouse for normal growth.

### *V dahliae* recovery assay

To determine the effects of a *V. dahliae* infection on cotton, we collected the stems and roots of the infected plants to analyse the fungal recovery potential. The samples were cut into many fragments and placed on PDA in plates, which were incubated at 25°C. After 5 days, the number of fragments with fungal hypha was recorded.

### Disease index (DI) analysis

The DI is an important parameter to assess plant resistance. The DI is calculated according using the following formula: DI = [(Σ disease grades × number of infected plants)/(total checked plants×4)] × 100. Seedlings were classified into five grades (grade 0, 1, 2, 3, and 4) based on the disease severity after *V. dahliae* infection as described by Wang *et al.* (2004).

## Results

### *V. dahliae* colonization and growth in the root interiors

In our previous studies, cotton plants started to show disease symptoms 15-18 days after inoculation with *V. dahliae*, including yellow leaves, defoliation and stunted growth. However, before the presence of the disease symptoms, it is unclear how the plants resist colonization and the upward dispersion of the fungal pathogen in the interior of plants (primarily xylem vessels). To investigate the colonization of the pathogen and its spread in the xylem vessels, root samples from the seedlings inoculated with *V. dahliae* for 1, 4, 7, 10 and 13 days were first treated by ultrasound to remove the fungal hyphae and spores on their surface, and the fungal DNA levels were examined by qPCR. The DNA of *V. dahliae* was barely detectable at 1 and 4 days post inoculation (dpi), indicating that few pathogens enter the root interior. While at 7 dpi, a few fungal DNA molecules were monitored in the roots, a value of 1.03×10^−4^ compared to the cotton DNA copies; at 10 and 13 dpi, relative DNA copies of the fungal pathogen were approximately 1.25×10^−3^ and 1.52×10^−3^, respectively, suggesting that the *V. dahlia* hyphae and conidia had located in the xylem vessels (Figure 1A). To investigate when the fungus colonizes in the interior of roots in more detail, a fungal recovery assay of the treated root fragments was conducted as a parallel experiment. Consistent with the results of the fungal DNA analysis, fungi were not observed around the root sections at 1 and 4 dpi, while approximately 5% of the root fragments showed fungi growth at 7 dpi, and a few fragments at 10 and 13 dpi demonstrated recoverable growth of the fungi (Figure 1B and 1C). These results suggested that *V. dahliae* had colonized in the inside of the roots through root-dipped inoculation after 7 dpi. To evaluate the interaction of the plant with the pathogen associated with miRNA regulation function in the root vascular tissue, the fungal-treated roots at the two time-points, 7 and 10 days, were chosen for sRNA high-throughput 361 sequencing analysis.

**Figure 1.**
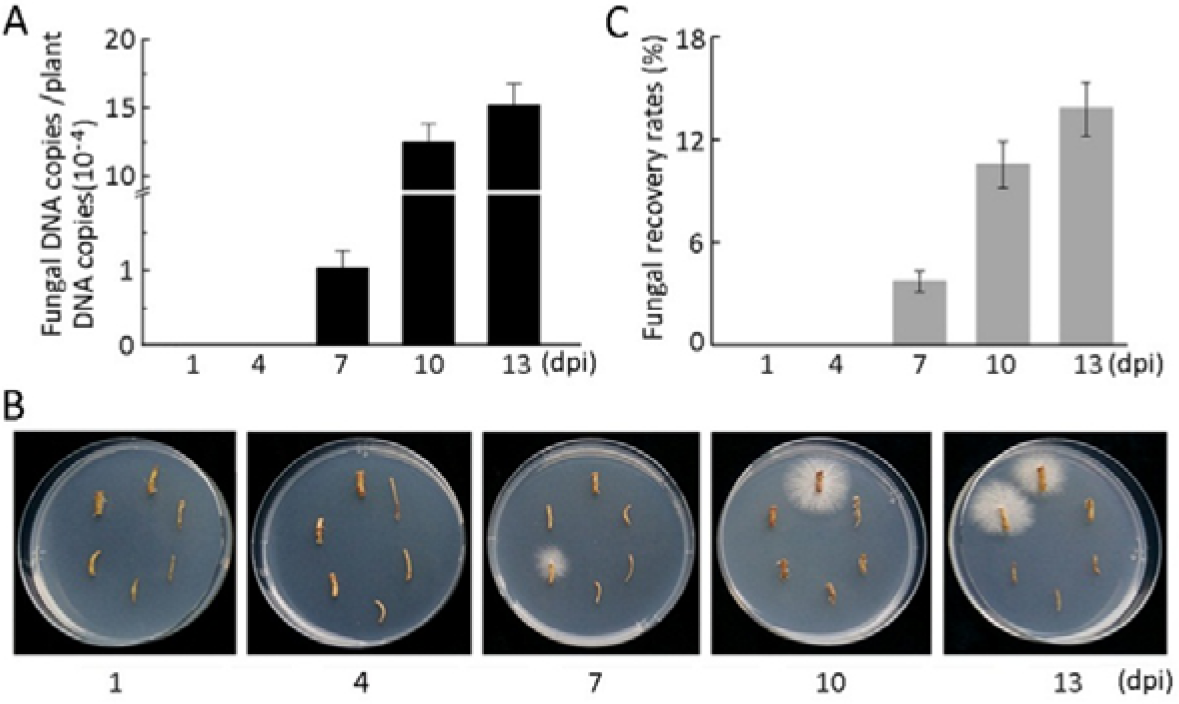
The time course of *V. dahliae* colonization and growth in the interior of cotton roots. (A) Fungal DNA copies/plant DNA copies in roots inoculated with *V. dahliae.* (B) Fungal recovery growth from the root fragments placed on PDA media at different time points of fungal inoculation. Photos were taken at 5 days after plating. (C) *V. dahliae* recovery rate of the root fragments (n=90). Error bars represent the SD of three biological replicates. dpi represents day post inoculation.

### High-throughput sequencing of sRNA

To characterize the sRNA profiles in the cotton plants challenged by *V. dahliae* infection, three sRNA libraries were constructed by using total RNA isolated from the root samples of seedlings treated for 7 and 10 days and the mixed mock treatment of 7 and 10 days (the control, CK) with two replicates for each treatment. The three libraries were sequenced with an Illumina HiSeq 2500, and a schematic flow of the sequencing and data analysis strategy is shown in Figure S1. As shown in Table 1, the three libraries for 7 and 10 dpi and the control generated more than 20 million clean reads, 20221992, 35104139 and 34150616, respectively. Through annotation analysis with the Silva, GtRNAdb, Rfam and Repbase databases, the sRNAs were grouped into several classes: repeat bases, rRNA, tRNA, snoRNA, and unannotated sRNA (Table 1). Before analysing the miRNA, the unannotated sRNA was mapped to the *G. hirsutum* cv TM-1 genome. A total of 3775953, 3944094 and 4889704 reads in the 7-and 10-dpi and control libraries, respectively, were successfully matched back to the AD genome of *G. hirsutum*, respectively (Table 1). Although the total reads of the 7 dpi sample were less than those of the 10 dpi and the control, the numbers of mapped unannotated reads containing miRNA were similar among the three libraries.

**Table 1.**
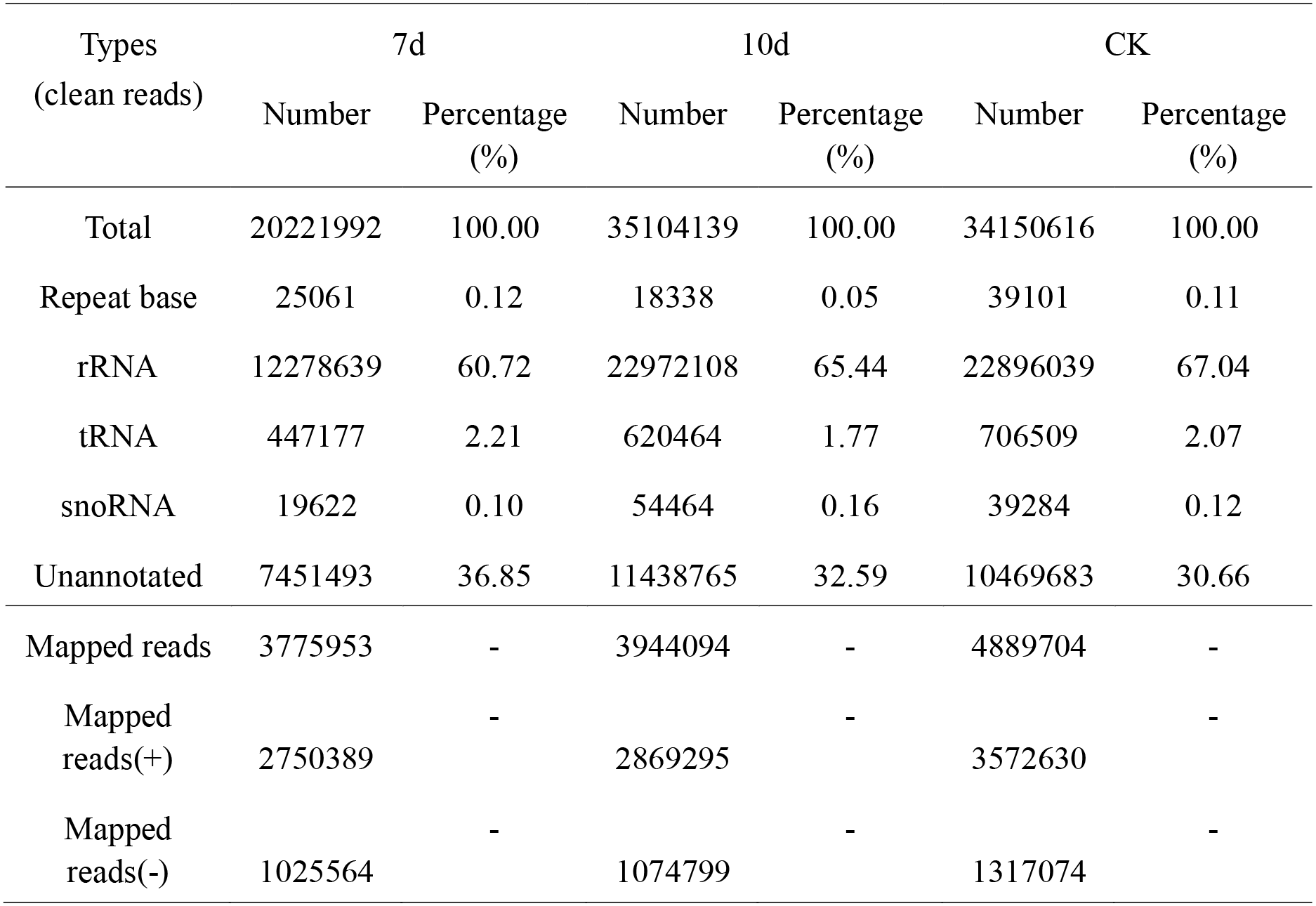
Cotton sRNA categorization and unannotated reads matched to the genome.

| Types<br>(clean reads) | 7d |  | 10d |  | CK |  |
| --- | --- | --- | --- | --- | --- | --- |
|  | Number | Percentage<br>(%) | Number | Percentage<br>(%) | Number | Percentage<br>(%) |
| Total | 20221992 | 100.00 | 35104139 | 100.00 | 34150616 | 100.00 |
| Repeat base | 25061 | 0.12 | 18338 | 0.05 | 39101 | 0.11 |
| rRNA | 12278639 | 60.72 | 22972108 | 65.44 | 22896039 | 67.04 |
| tRNA | 447177 | 2.21 | 620464 | 1.77 | 706509 | 2.07 |
| snoRNA | 19622 | 0.10 | 54464 | 0.16 | 39284 | 0.12 |
| Unannotated | 7451493 | 36.85 | 11438765 | 32.59 | 10469683 | 30.66 |
| Mapped reads | 3775953 | - | 3944094 | - | 4889704 | - |
| Mapped<br>reads(+) | 2750389 | - | 2869295 | - | 3572630 | - |
| Mapped<br>reads(-) | 1025564 | - | 1074799 | - | 1317074 | - |

Notes: rRNA, ribosomal RNA; tRNA, transfer RNA; snoRNA, small nucleolar RNAs; Mapped reads (+) and Mapped reads (-), the number of clean reads matched on the positive and the negative chains from the cotton genome.

To further ensure the specificity and commonality of the sRNA in the three libraries, the unique reads calculated in the 7- and 10-dpi and the control sRNA libraries were 4686834, 6144606, and 6024129, respectively. There were specific and common sequence types shown in the comparison of the two libraries, such as 3581183 and 4918478 specific unique reads and 1105651 common unique reads in the 7 dpi vs the control libraries, 5130747 and 5013270 specific unique reads and 1010859 common unique reads in the 10 dpi vs the control libraries, and 3815094 and 5269866 specific unique reads and 871740 common unique reads in the 7 dpi vs 10 dpi libraries (Figure 2A).

**Figure 2.**
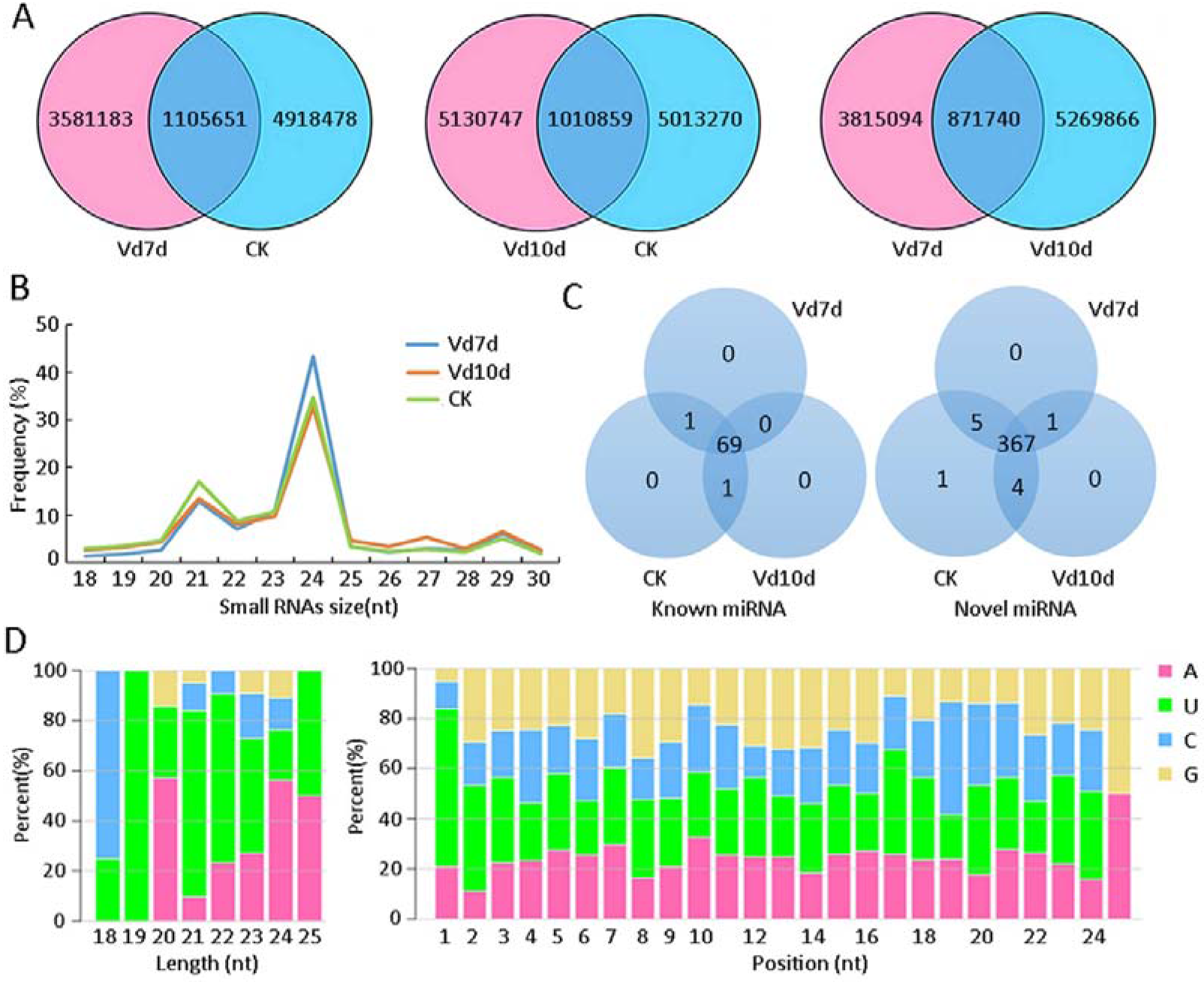
sRNA and miRNA analyses of the 7- and 10-dpi libraries and the control. (A) Venn diagram for special and common unique reads between the libraries. (B) Size distribution of the matched sRNA reads in cotton. (C) Venn diagram for special and common known and novel miRNAs among the three libraries. (D) miRNA first nucleotide bias (left panel) and miRNA nucleotide bias at each position (right panel) among the three sRNA libraries. Vd7d and Vd10d, represent 7 and 10 days after *V. dahliae* infection. CK, mixed mock treatment samples at 7 and 10 days.

To investigate the size distribution of all the sequences, the sequences between 18 and 30 nt were determined in the number of matched unannotated reads. The size distribution for the matched reads was similar through observation of the three libraries, in which the 24 nt reads accounted for the majority, 43.41%, 32.93% and 34.77% for 7 dpi, 10 dpi and the control, respectively, followed by 21 nt reads, accounting for 12.86%, 13.41% and 16.99%, respectively (Figure 2B). The results of the sRNA abundance and size in cotton were consistent with previous reports in cotton (Wang *et al.*, 2016) and consistent with the results reported in *Arabidopsis thaliana* (Rajagopalan *et al.*, 2006), *Oryza sativa* (Wei *et al.*, 2011), and *Glycine max* (Song *et al.*, 2011), suggesting that the sRNAs in plants are mainly composed of 21 and 24 nt reads.

### Identification of the miRNAs

By using miRDeep2 analysis, we screened the unannotated sRNA sequences to identify miRNAs according to the criteria for the selection of a length of at least 18 nt and a maximum of two mismatches compared to all known plant miRNA sequences in the three libraries. After removing the repeat sequences, 71 annotated known miRNAs belonging to 46 miRNA families were identified; out of these, 70, 70 and 71 were from the 7- and 10-d treated roots and the control roots, respectively (Supplementary Table S1). Of the 71 miRNAs across the three libraries, 69 miRNAs were commonly present in the three libraries, while ghr-miR7497 and ghr-miR399e were absent in the 7- and 10-dpi libraries, respectively (Figure 3C). Each of the 46 miRNA families contained 1 to 4 members. The three families, MIR156, MIR2949 and MIR482, possessed 4 members, while there were 30 other miRNA families with only one member (Figure S2). As shown in Supplementary Table S2, the expression levels of the miRNAs ranged widely from tens of thousands of sequence reads to fewer than 100. The MIR166 had the most abundant expression and reached over 10000 TPM clean reads, which are highly conserved in mosses, eudicots and monocots (Arazi *et al*., 2005; Barik *et al*., 2014; Guo *et al.*, 2017; Shi *et al*., 2017; Yip *et al.*, 2016). In addition, among the 46 miRNA families, 21 and 24 nt long miRNAs represented the majority in size, reaching 38.03% and 32.39%, respectively, followed by the 20 nt long miRNAs (14.08%) (Supplementary Table S3).

**Figure 3.**
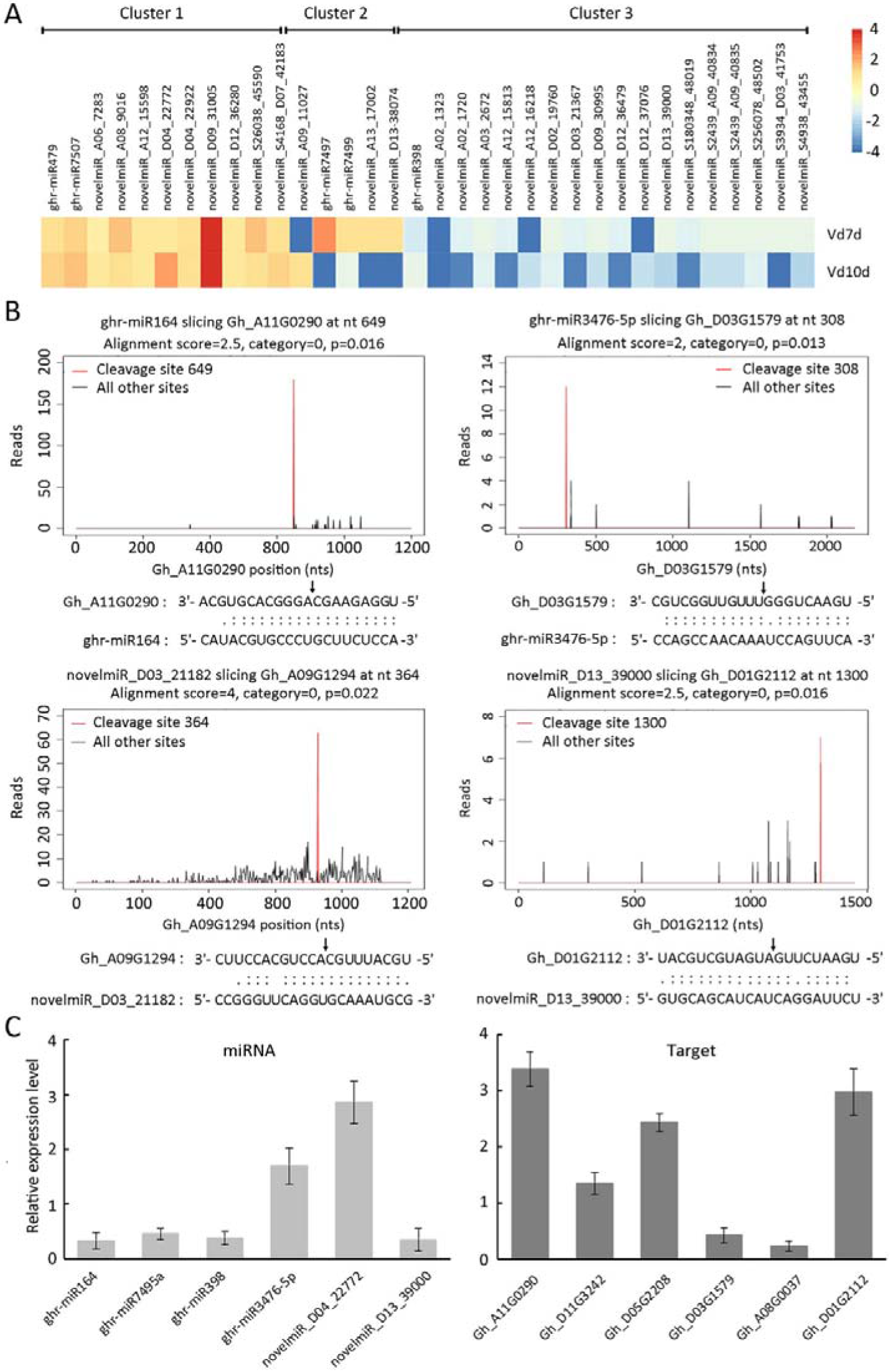
Analyses of different expression of the miRNAs and target gene identification as well as expression authenticity. (A) Heatmaps of differently expressed miRNA in the 7- and 10-d treated libraries compared to the control. The colour bar represents the relative signal intensity values from red (upregulated) to blue (downregulated), indicating a range of [4,−4]. (B) Cotton miRNA and target alignment and its T-plot validated by degradome sequencing. The T-plots indicate the distribution of the degradome tags along the full length of the target mRNA sequencing. The black arrows indicate the cleavage sites of the target genes. (C) Expression profiles of miRNAs and corresponding targets after *V. dahliae* was inoculated by qPCR. miRNAs and their corresponding targets detected from the roots infected with *V. dahliae* and mock-treated control at 10 dpi, respectively. Error bars represent the SD of three biological replicates.

To identify novel miRNAs, the unannotated sRNAs, which could be mapped to the cotton AD genome excluding the known miRNA, were screened using miRDeep2 software. A total of 378 unknown sRNA sequences were supposed to be novel miRNAs with high confidence in the three libraries. There were 373, 372 and 377 novel miRNAs in the 7- and 10-dpi libraries and the control library, respectively. Among these 378 novel miRNAs, approximately 367 were common across all three libraries, and only one novel specific miRNA (novel miR_A02_1323) was detected in the control library. The novel miR_D09_31005 was only found in the two treated libraries (Figure 2C).

In this study, nucleotide bias at positions in the total miRNAs was analysed to understand the miRNA sequence law. The results demonstrated that the first nucleotide of the miRNAs exhibited a preference for uracil (U) (Figure 2D, left panel), consistent with the results from many species possibly due to miRNA sequence conservation. In addition, nucleotide bias at each position is also shown in Figure 2D (right panel) consistent with other plants (Mi *et al.*, 2008).

### miRNA expression response to *V. dahliae* infection and the corresponding target prediction

The identified miRNAs with more than 5 TPM expression levels were chosen for an analysis of differential expression among the 7- and 10-d treated roots and the control. As shown in Supplementary Table S4, 28 out of 71 (39.44%) known miRNAs and 148 out of 378 (39.15%) novel miRNAs were differentially expressed in the two pathogen-induced libraries compared to the mock-treated control (absolute value of log2 ratio ≥ 1). Among all the differentially expressed miRNAs, 5 and 29 differentially expressed miRNAs, respectively, from the known and novel miRNAs were found in both the 7- and 10-d treated roots compared to the control (Figure 3A). Twenty-nine miRNAs showed significant upregulation or down regulation of expression in both treated libraries (Cluster 1 and 3); 4 miRNAs exhibited upregulated expression in the 7-d treated roots and downregulation in the 10-d treated roots, and only one miRNA showed a contrasting trend (Cluster 2).

To further investigate the function of differentially expressed miRNAs, the corresponding target genes were predicted, and GO enrichment was performed. According to TargetFinder software and the GO classifications, the 405 target genes of the differentially expressed miRNAs were predicted and associated with many GO terms in the 7- and 10-d libraries compared to the control, primarily including binding and oxidoreductase activity (Supplementary Table S5 and S6).

### Identification and functional enrichment of the miRNA targets by degradome combined with sRNA sequencing

To identify the target genes from a total of 449 miRNAs, degradome sequencing through the next-generation deep sequencing technique was performed. A total of 178 target genes of the miRNAs in the cotton root RNA library was identified. Twenty-five of the 71 known miRNAs regulated 75 target transcripts, and 172 of the 378 novel miRNAs possessed 142 target genes (Supplementary Table S7 and Table S8). miRNAs were found to be able to target various numbers of genes with a range of 1 to 16, of which novelmiR_D05_23410 targeted the highest number of genes, reaching 16 different genes (Supplementary Table S8). As shown in Figure 3B, examples of some confirmed defence-related miRNA targets as ‘target plots’ (T-plots) were drawn, which describe the cleavage sites of the target sequences by the action of different miRNAs.

Among the 178 target genes identified from degradome sequencing confirmed by sRNA sequencing, 40 target genes were associated with the differentially expressed miRNAs. According to the GO classifications, the 40 target genes of the differentially expressed miRNAs predominantly participated in 61 biological process categories, 53 molecular function categories and 16 cellular component categories (Supplementary Table S9). Most specific GO classifications showed that the target genes were involved in the response to oxidation-reduction process, the response to biotic and abiotic stress, and binding (Supplementary Table S9). The KEGG analysis classified 11 different expression miRNA targets into 10 pathways, and the significantly enriched pathways included terpenoid backbone biosynthesis, carotenoid biosynthesis and spliceosome (Supplementary Table S10).

### Expression authenticity of miRNAs and the corresponding target genes using qPCR

To further confirm the authenticity of the sRNA high-throughput sequencing and identify targets by degradome sequencing, the expression abundance of the miRNAs and their corresponding target genes was tested by qPCR. Six different expression level miRNAs (4 known miRNAs and 2 novel miRNAs) and their corresponding target genes were chosen to analyse the expression levels in 10-d treated roots and the control. The expression levels of 4 miRNAs, ghr-miR164, ghr-miR3476-5, ghr-miR398 and novelmiR_D13_39000, decreased remarkably compared to the control, while the expression levels of ghr-miR7495a and novelmiR_D04_22772 increased significantly, similar to the results of the sRNA sequencing (Figure 3C and Supplementary Table S4). The expression levels of six corresponding target genes compared to the control showed contrasting trends with the miRNAs, indicating that the six miRNAs negatively regulated the mRNA levels of the corresponding target genes (Figure 3C).

### Expression regulation of *GhNAC100* by a ghr-miR164 post-transcriptional process

Based on the degradome and sRNA sequencing, we selected ghr-miR164 to further evaluate the regulatory functions of the miRNA-target modules in the resistance of the plant to the fungus. The ghr-miR164 target gene was identified as Gh_A11G0290, shown to be an NAC domain-containing protein 100-like according to the BLAST data. A phylogenetic tree shows that cotton NAC 100-like is clustered with AtNAC100 (57.36% identification) and was designated GhNAC100 (Figure 4A). The results of qPCR analysis showed that the ghr-miR164 expression level decreased approximately 65% in the 10-d treated roots compared to the control consistent with the sRNA sequencing data, while *GhNAC100* was upregulated in the treated roots, approximately 3.4-fold higher than in the control roots (Figure 3C).

In our degradome sequencing data, ghr-miR164 matched the target gene *GhNAC100*, and the cleavage site was identified in a matched sequence as shown in the T-plots (Figure 3B). The cleavage site is located at the 649 nt of *GhNAC100* mRNA, which was cleaved between the G and C bond (Figure 3B). To confirm this cleavage site, two specific forward primers were designed, which were located on both sides of cleavage site, respectively (Figure 4B), and qPCR analysis was conducted. As shown in Figure 4B, the amounts of the FD fragment downstream of the cleavage site were approximately 1.7-fold higher than the FU fragment containing the cleavage site.

**Figure 4.**
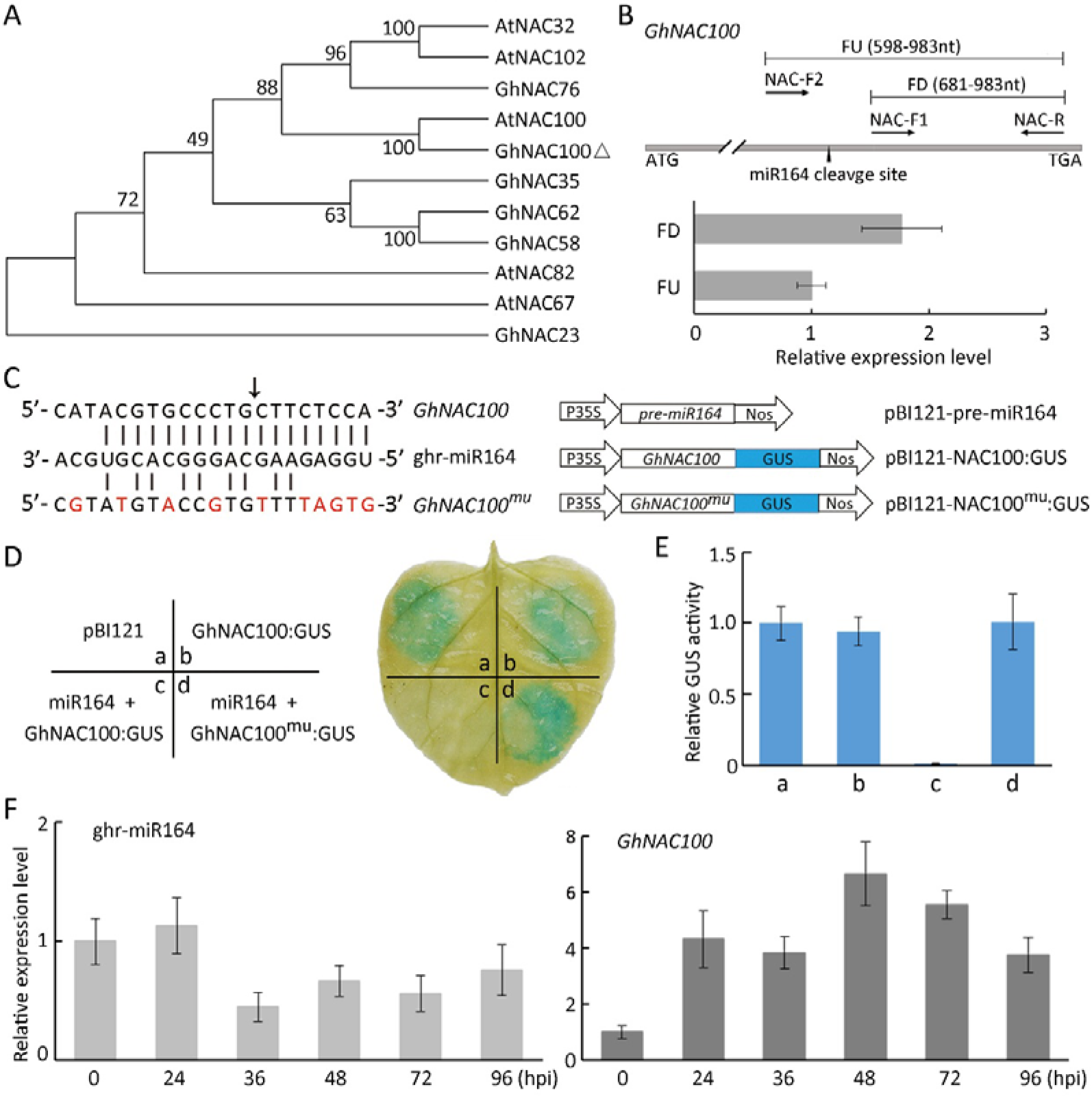
ghr-miR164 regulates *GhNAC100* expression by a post-transcriptional process. (A) Phylogenetic tree analysis of GhNACs and AtNACs. The Neighbour-joining method of MEGA (version 5.2) was used. Bootstrap analyses were computed with 1000 replicates. Accession numbers are: AtNAC32 (AT1G77450), AtNAC102 (AT5G63790), GhNAC76 (AHJ79217.1), AtNAC100 (AT5G61430), GhNAC35 (AHJ79176.1), GhNAC62 (AHJ79203.1), GhNAC58 (AHJ79199.1), AtNAC82 (AT5G09330), AtNAC67 (AT4G01520), and GhNAC23 (AHJ79164.1). (B) Primer design outline and qPCR analysis of the *GhNAC100* mRNA transcripts associated with the ghr-miR164 cleavage. (C) Construction of the effector and reporter vectors. Red letters represent mutated bases. The black arrow indicates the cleavage site of *GhNAC100.* (D) GUS staining of infiltrated sites of the leaf with different vectors. (E) Quantitative analysis of GUS activity from (D) with the 4-MU assay. (F) Accumulation of ghr-miR164 and *GhNAC100* in the time course of cotton roots infected with *V. dahliae.* Error bars represent the SD of three biological replicates.

To further verify the ghr-miR164 function in cleaving its target sequence *in vivo*, a GhNAC100:GUS reporter fusion protein was analysed by the *Agrobacterium tumefaciens-mediated* co-transformation technology in *Nicotiana benthamiana.* The precursor of ghr-miR164 was isolated and inserted into a plant expression vector driven by the CamV35S promoter, resulting in the construction of vector pBI121-pre-miR164 as an effector. The *GhNAC100*-encoding sequence and its corresponding mutant sequence were respectively fused into the upstream of the *GUS* gene in the plant expression vector pBI121, generating pBI121-GhNAC100:GUS and pBI121-GhNAC100^mu^:GUS as reporters (Figure 4C). As shown in Figure 4D, the leaves injected with GV3101 only containing pBI121 or pBI121-GhNAC100:GUS exhibited a similar blue intensity in the infiltrated site by GUS histochemical staining. When the leaves were infiltrated with equally mixed GV3101 cells containing pBI121-pre-miR164 or pBI121-GhNAC100:GUS, the blue spot was absent at the injected site, while the leaves co-infiltrated with mixed GV1301 cells containing pBI121-pre-miR164 or pBI121-GhNAC100^mu^:GUS showed a similar blue intensity to those only infiltrated with pBI121-GhNAC100:GUS (Figure 4D). Compatible with GUS staining, a quantitative assay of the GUS activity showed similar results in the extracted total protein from the infiltrated sites of the leaf as indicated through 4-MU analysis (Figure 4E). The result of the GUS fusion protein reporter showed that ghr-miR164 could cleave *GhNAC100* by a post-transcriptional process *in vivo*.

To explore the role of the ghr-miR164-GhNAC100 module in the response of the plant to fungal infection, qPCR analysis was performed to measure the time course of the pathogen-responsive expression profile of ghr-miR164 and *GhNAC100.* The results showed that the accumulation of ghr-miR164 decreased in the roots and reached a minimum level at 36 hpi (Figure 4F). In contrast, the *GhNAC100* transcript level increased in the roots of plants challenged with *V. dahliae* and reached a maximal level at 48 hpi (Figure 4F). These data suggest that the ghr-miR164 content negatively regulated the *GhNAC100* expression level, participating in the plants inoculated with *V. dahliae.*

### ghr-miR164 silencing improves plant resistance to *V. dahliae*

To determine the function of ghr-miR164 in plant defence, miRNA target mimicry technology was employed, which has been successfully used to suppress miRNA accumulation *in vivo* (Sha *et al*., 2014; Yan *et al*., 2012). We used the virus-based microRNA silencing (VbMS) strategy to generate the ghr-miR164-silenced plants. The pTRV2e-STTM164 vector, which contains two imperfect binding sites for ghr-miR164 separated by a 48 nt spacer, was constructed. The cotton *phytoene desaturase (GhPDS)* gene, a positive control, was well silenced resulting in a photobleaching phenotype (Figure S3), indicating that the TRV VIGS system is feasible in the cotton plant. Compared with the negative control plants inoculated with the empty vector (TRV:00, as the control), the abundance of ghr-miR164 transcripts was significantly reduced by approximately 50% in the *TRV:STTM164* cotton plants (Figure 5A). The *GhNAC100* expression level was also tested in these infected plants, showing an increase of approximately 2.5-fold compared to the *TRV:00* plants (Figure 5A). These data suggested that we successfully knocked down the ghr-miR164 expression in the *TRV:STTM164* plants by overexpressing STTM using the VbMS. *TRV:STTM164* plants and the control were infected with 10^6^ *V. dahliae* conidia through root-dipped inoculation. After 23 dpi, the *TRV:00* plants showed normal disease symptoms with wilting, yellowing leaves and stunted growth, while the *TRV:STTM164* plants exhibited obvious resistance to this fungus (Figure 5B). The DI value in the *TRV:STTM164* plants was significantly lower than that of the control plants, showing value 38, while the control showed 53 (Figure 5C). To examine the extent of the *V. dahliae* colonization in the infected stems, a fungal recovery assay was performed. There were fewer stem sections that provided fungal colonies in the *TRV:STTM164* plants than those in the *TRV:00* plants (Figure 5D). Consistent with the fungal recovery assay, the fungal biomass in the ghr-miR164-silenced plants decreased significantly to approximately 0.3-fold of the control plants (Figure 5E).

**Figure 5.**
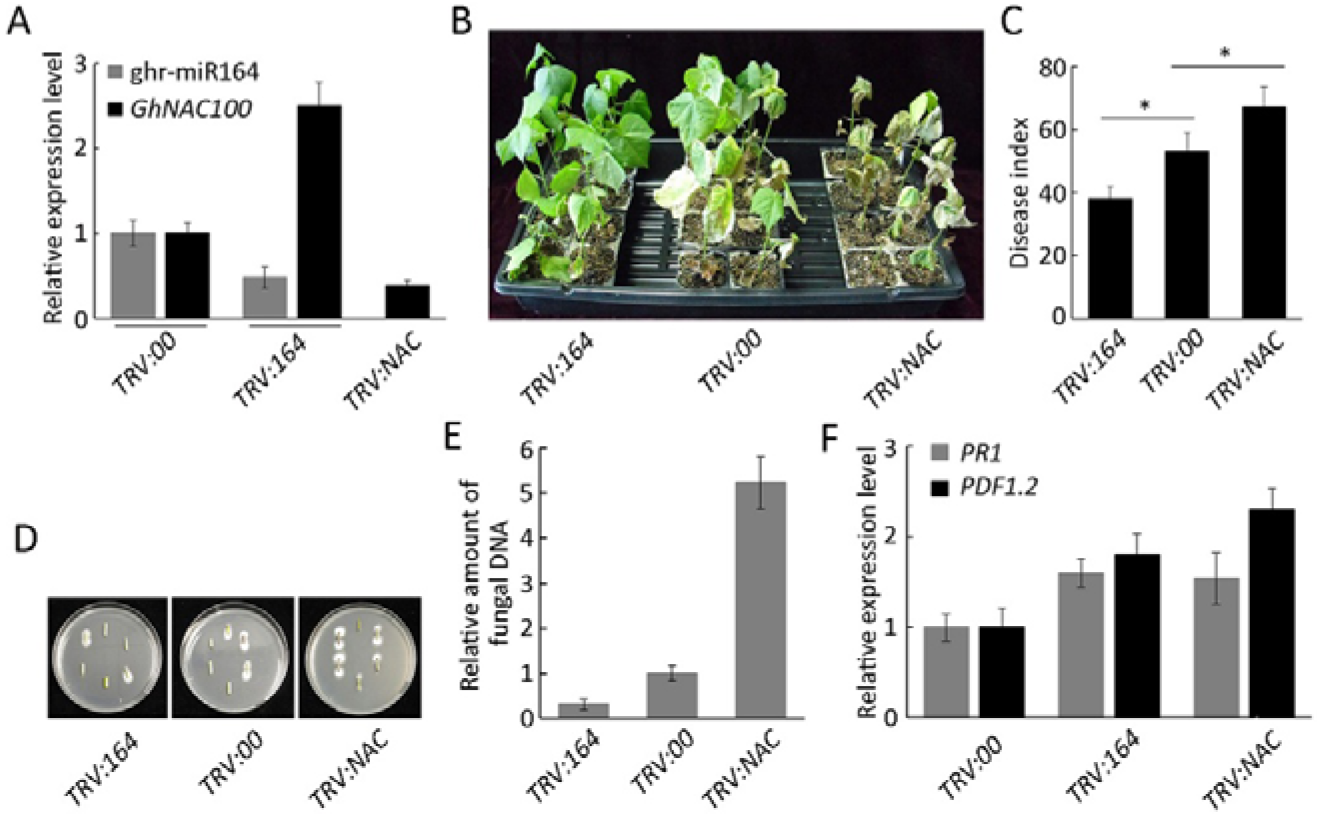
The functional dissection of the ghr-miR164-*GhNAC100* module in defence against V *dahliae*. (A) Relative expression levels of ghr-miR164 and *GhNAC100* in the *TRV:STTM164* and *TRV:GhNAC100* compared to the *TRV:00* plants. (B) Disease symptom phenotypes of the ghr-miR164-silenced and *GhNAC100*-silenced plants inoculated with *V. dahliae.* (C) Disease index of the silenced plants at 23 dpi. Significant differences were determined using Student’s *t*-test; an asterisk indicates *P* < 0.05. (D) Fungal recovery assay. The experiment was performed using the stem sections from cotton plants at 23 dpi placed on PDA media. Photos were taken 5 days after plating. (E) The levels of the *V. dahliae* biomass in the infested stems by qPCR. Error bars represent the SD of three biological replicates. *TRV:164* and *TRVNAC* represent *TRV:STTM164* and *TRV:GhNAC100*, respectively.

### The ghr-miR164-GhNAC100 module regulates plant defence to *V. dahliae*

To clarify the roles of the ghr-miR164-GhNAC100 module in the resistance of the plant to *V. dahliae*, the *GhNAC100* gene was knocked down by a tobacco rattle virus (TRV)-mediated VIGS system. When the *PDS*-silenced plant leaves became chlorotic, we started to examine the *GhNAC100* expression levels of the plants injected by Agrobacterium containing the *TRV:GhNAC100* virus vector. According to the qPCR analysis, *GhNAC100* accumulation in the leaves infected with *TRV:NAC100* significantly decreased to approximately 61% of the plants infected with the *TRV:00* (Figure 5A). To evaluate the GhNAC100 function in the resistance to this fungus, the GhNAC100-silenced plants and the control were infected with *V. dahliae* through root-dipped inoculation. After 23 dpi, the *TRV:NAC100* plants showed more serious disease symptoms than the control plants with obvious necrotic and wilting leaves and stunted growth (Figure 5B). The DI value in the *GhNAC100-silenced* plants was significantly higher than that in the control (Figure 5C). A fungal recovery assay was performed to examine the extent of the *V. dahliae* colonization in the infected stem of the treated plants. The results showed that there were more fungal colonies in the *TRV:GhNAC100* plants than in the *TRV:00* plants (Figure 5D). Consistent with these results, the fungal biomass in the *GhNAC100-silenced* plants increased significantly, 5.2-fold higher compared to the control plants. The results showed that *GhNAC100* is possibly a positive regulator to increase plant resistance to *V dahliae* (Figure 5E).

To investigate whether the regulation of the ghr-miR164-GhNAC100 module in plant defence was associated with the salicylic acid (SA) and jasmonic acid (JA) signalling pathways, the expression levels of both defence-related genes, *PR1* (SA signalling pathway) and *PDF1.2* (JA signalling pathway), were monitored in the *TRV:STTM164* and *TRV:GhNAC100* plants. As shown in Figure 5F, the *PR1* expression level increased remarkably in ghr-miR164-silenced plants compared to the wild-type plants, while it significantly decreased in the *GhNAC100*-silenced plants. Interestingly, the transcript levels of *PDF1.2* showed similar results to those of *PR1* in the *TRV:STTM164* and *TRV:GhNAC100* plants. The results indicated that the ghr-miR164-GhNAC100 module in plant defence may be involved in both the SA and JA signalling pathways.

## Discussion

miRNAs participate in plant resistance to pathogens through post-transcriptional regulation of the expression level of the target genes. The response of the cotton plants to *V. dahliae* infestation may be divided into early and later stages, including the pathogen localizing in the surface of roots and entering their interior. Previous reports involving sRNA sequencing focused only on the early response of the plant to *V. dahliae* infection, which in general reacts before 48 hours after inoculation. However, there are few reports about the later response of the plant to this fungus after 4 days. In this study, we constructed 7- and 10-d infected root sRNA libraries with *V. dahliae* and identified many known and novel miRNAs. Differentially expressed miRNAs of the 7- and 10-day root-infected libraries were analysed compared to the mock control combined with degradome sequencing. The results showed that 71 known miRNAs and 378 novel miRNAs were identified from 7- and 10-day root-infected and control libraries, and 34 differentially expressed miRNAs in both 7- and 10-d infected roots were analysed compared to the control. More importantly, a ghr-164-GhNAC100 module was selected to represent the miRNAs and corresponding targets to perform functional dissection of plant defence against *V dahliae*.

We identified 378 novel miRNAs from three sRNA libraries of roots inoculated with *V. dahliae* and the mock-treated control at 7- and 10-dpi when the fungus has entered the inner root tissues. However, previous studies showed that significantly fewer novel miRNAs were identified at 12, 24, 36 or 48 hours after inoculation (He *et al*., 2014; Yin *et al.*, 2012; Zhang *et al.*, 2015a). For instance, 14 novel miRNAs were identified from *Verticillium-inoculated* cotton roots at 12 and 24 hours (Yin *et al*., 2012), 37 novel miRNAs were identified from two sRNA libraries of the cotton seedlings inoculated with *V. dahliae* at 24 and 48 hours (He *et al.*, 2014), and 58 novel miRNAs were identified from sRNA sequencing of the cotton roots inoculated with *V. dahliae* at 24 hours (Zhang *et al*., 2015a). Thus, in this study, more novel miRNAs were identified, which may participate in the resistance of the plant to *V. dahliae* at the later stages of fungal infection.

At 7 and 10 days after inoculation, the plants should have a stronger response to fungal infection, possibly because *V. dahliae* has localized in the xylem vessels, unlike the pathogen surface-induced response at the early stage of inoculation. According to the GO analysis, the target genes of the differentially expressed miRNAs predominantly participated in many GO terms, including the oxidation-reduction process and stress response (Supplementary Table S9). KEGG analysis classified 11 miRNA targeting 10 pathways, and the significantly enriched pathways include terpenoid backbone biosynthesis, carotenoid biosynthesis and the spliceosome (Supplementary Table S10). However, in the literature associated with fungal surface-induced miRNAs, there is no data on the GO and KEGG analyses of target genes (He *et al*., 2014; Yin *et al.*, 2012; Zhang *et al*., 2015a). These results suggested that the internal induced response of the plant by *V. dahliae* infection may be stronger to participate in the resistance by modulating miRNA expression to post-transcriptionally regulate the target gene.

Based on sRNA and degradome sequencing, we chose the ghr-miR164-GhNAC100 module as representative to analyse the function of the miRNAs coupling with their target genes in the resistance of the plant to *V dahliae.* Our results showed that ghr-miR164 silencing elevated the resistance of the plants to this fungus, consistent with the results of its target *GhNAC100* knockdown, which increased the susceptibility of the plant to the pathogen. Therefore, the ghr-miR164-GhNAC100 module participates in plant defence against *V. dahliae*. Currently, there were some reports that miR164 modulates plant resistance through post-transcriptional regulation of its target gene expression. For instance, Arabidopsis NAC4 promoted pathogen-induced cell death under negative regulation by microRNA164 (Lee *et al*., 2017). Feng *et al.* (2014) reported that TaNAC21/22 participated in the resistance of wheat plants to stripe rust regulated by tae-miR164. In summary, our results documented that miR164 participates in plant defence against pathogens by post-transcriptionally regulating the expression of the NAC transcriptional factor.

In addition, the NAC is negatively regulated by miR164 through mRNA cleavage to participate in development excluding defence (Baker *et al.*, 2005; Sieber *et al.*, 2007). In Arabidopsis, miR164 targets the transcripts of six *NAC* genes and prevents organ boundary enlargement and the formation of extra petals during flower development (Laufs *et al*., 2004; Mallory *et al*., 2004). ORE1, an NAC transcription factor, is involved in leaf cell death through miRNA164 regulation (Kim *et al*., 2009). In maize (*Zea mays* L.), miR164-directed cleavage of *ZmNAC1* confers lateral root development (Li *et al.*, 2012). However, our study was focused on the plant resistance through VIGS methods. In addition, the phenotype of the gene-silenced plants was shown in the early stage of development, the seedling. Of course, it would be interesting to investigate whether ghr-miR164 affects plant development in the future.

Previous reports involving the sRNA sequencing of cotton plants in response to *V. dahliae* focused on the early induction stage, typically 12-48 hours after inoculation when the fungus localized on the surface of the roots. We acquired the sRNA profiles of the plant response to *V. dahlia*, which had localized in the interior root tissues, a later induction stage. We identified 71 known miRNAs and 378 novel miRNAs from 7- and 10-d *V. dahliae*-infected libraries and the control library and investigated their target categories using GO and KEGG analyses. Thirty-four of these miRNAs showed significantly different expression in the two infected libraries compared to the control. More importantly, according to the degradome and sRNA sequencing, we selected the ghr-miR164-GhNAC100 module as representative to evaluate the function of the miRNAs in the response of the plant to the fungus through post-transcriptional regulation of the expression level of the target genes. The results showed that ghr-miR164-GhNAC100 participates in cotton plant resistance to *V. dahliae.*

## Supplementary data

**Table S1.** Summary of known miRNAs.

**Table S2.** Read abundance of known miRNAs.

**Table S3.** Size distribution of known miRNAs.

**Table S4.** Differentially expressed miRNAs identified in *V. dahliae-infected* cotton compared to control.

**Table S5.** GO enrichment analysis of the predicted targets of differentially expressed known and novel miRNAs in Vd7d library vs the control library.

**Table S6.** GO enrichment analysis of the predicted targets of differentially expressed known and novel miRNAs in Vd10d library vs the control library.

**Table S7.** List of target genes from degradome sequencing combining to differentially expressed known miRNAs.

**Table S8.** List of target genes from degradome sequencing combining to differentially expressed novel miRNAs.

**Table S9.** GO enrichment analysis of the targets of differentially expressed known and novel miRNAs based on degradome sequencing.

**Table S10.** KEGG enrichment analysis of the targets of differentially expressed known and novel miRNAs based on degradome sequencing.

**Table S11.** The primer sequences used in this study.

**Figure S1.** Schematic representation of analysis pipeline.

**Figure S2.** Member numbers of the known miRNA families

**Figure S3.** The phenotype of the GhPDS-silenced plants after VIGS treatment.

## Acknowledgements

This work was supported by the National Natural Science Foundation of China (31771848 and 31471544) and sponsored by the National Transgenic Major Programmes of China (2016ZX08005-003-002 and 2018ZX0800901B).

## Abbreviations

miRNA: microRNA
sRNA: small RNA
NAC: **n**o apical meristem **A**rabidopsis transcription activation factor and **c**up-shaped cotyledon
qPCR: real-time quantitative polymerase chain reaction
dpi: days post inoculation
VIGS: virus-induced gene silencing
TRV: tobacco rattle virus
PDS: Phytoene desaturase
DI: disease index
STTM: small tandem target mimic
JA: jasmonic acid
SA: salicylic acid

## References

Allen E, Xie Z, Gustafson AM, Carrington JC. 2005. microRNA-directed phasing during trans-acting siRNA biogenesis in plants. Cell 121, 207–221.

Anders S, Huber W. 2010. Differential expression analysis for sequence count data. Genome Biology 11, R106.

Arazi T, Talmor-Neiman M, Stav R, Riese M, Huijser P, Baulcombe DC. 2005. Cloning and characterization of micro-RNAs from moss. The Plant Journal 43, 837–848.

Ashburner M, Ball CA, Blake JA, et al. 2000. Gene Ontology: tool for the unification of biology. Nature Genetics 25, 25.

Baker CC, Sieber P, Wellmer F, Meyerowitz EM. 2005. The early extra petals 1 mutant uncovers a role for microRNA miR164c in regulating petal number in Arabidopsis. Current Biology 15, 303–315.

Barik S, SarkarDas S, Singh A, Gautam V, Kumar P, Majee M, Sarkar AK. 2014. Phylogenetic analysis reveals conservation and diversification of microRNA166 genes among diverse plant species. Genomics 103, 114–121.

Bazzini AA, Almasia NI, Manacorda CA, et al. 2009. Virus infection elevates transcriptional activity of miR164a promoter in plants. BMC Plant Biology 9, 152.

Bazzini AA, Hopp HE, Beachy RN, Asurmendi S. 2007. Infection and coaccumulation of tobacco mosaic virus proteins alter microRNA levels, correlating with symptom and plant development. Proceedings of the National Academy of Sciences of the United States of America 104, 12157–12162.

Bejaranoalcazar J, Blancolopez MA, Melerovara JM, Jimenezdiaz RM. 1997. The influence of verticillium wilt epidemics on cotton yield in southern spain. Plant Pathology 46, 168–178.

Bhat RG, Subbarao KV. 1999. Host range specificity in verticillium dahliae. Phytopathology 89, 1218–1225.

Fahlgren N, Howell MD, Kasschau KD, et al. 2007. High-throughput sequencing of Arabidopsis microRNAs: evidence for frequent birth and death of MIRNA genes. PLoS One 2, e219.

Feng H, Duan X, Zhang Q, Li X, Wang B, Huang L, Wang X, Kang Z. 2014. The target gene of tae-miR164, a novel NAC transcription factor from the NAM subfamily, negatively regulates resistance of wheat to stripe rust. Molecular Plant Pathology 15, 284–296.

Friedlander MR, Mackowiak SD, Li N, Chen W, Rajewsky N. 2012. miRDeep2 accurately identifies known and hundreds of novel microRNA genes in seven animal clades. Nucleic Acids Research 40, 37–52.

German MA, Pillay M, Jeong DH, et al. 2008. Global identification of microRNA-target RNA pairs by parallel analysis of RNA ends. Nature Biotechnology 26, 941–946.

Guo Y, Zhao S, Zhu C, Chang X, Yue C, Wang Z, Lin Y, Lai Z. 2017. Identification of drought-responsive miRNAs and physiological characterization of tea plant (Camellia sinensis L.) under drought stress. BMC Plant Biology 17, 211.

Hao J, Tu L, Hu H, Tan J, Deng F, Tang W, Nie Y, Zhang X. 2012. GbTCP, a cotton TCP transcription factor, confers fibre elongation and root hair development by a complex regulating system. Journal of Experimental Botany 63, 6267–6281.

He X, Sun Q, Jiang H, et al. 2014. Identification of novel micrornas in the verticillium wilt-resistant upland cotton variety kv-1 by high-throughput sequencing. Springerplus 3, 564.

Jefferson RA, Kavanagh TA, Bevan MW. 1987. GUS fusions: beta-glucuronidase as a sensitive and versatile gene fusion marker in higher plants. The EMBO Journal 6, 3901–3907.

Jia X, Ren L, Chen QJ, Li R, Tang G. 2009. UV-B-responsive microRNAs in Populus tremula. Journal of Plant Physiology 166, 2046–2057.

Jones-Rhoades MW, Bartel DP, Bartel B. 2006. MicroRNAs and their regulatory roles in plants. Annual Review of Plant Biology 57, 19–53.

Kanehisa M, Goto S, Kawashima S, Okuno Y, Hattori M. 2004. The KEGG resource for deciphering the genome. Nucleic Acids Research 32, D277–D280.

Khraiwesh B, Zhu JK, Zhu J. 2012. Role of miRNAs and siRNAs in biotic and abiotic stress responses of plants. Biochimica et Biophysica Acta 1819, 137–148.

Kim JH, Woo HR, Kim J, Lim PO, Lee IC, Choi SH, Hwang D, Nam HG. 2009. Trifurcate feed-forward regulation of age-dependent cell death involving miR164 in Arabidopsis. Science 323, 1053–1057.

Klosterman SJ, Atallah ZK, Vallad GE, Subbarao KV. 2009. Diversity, pathogenicity, and management of verticillium species. Annual Review of Phytopathology 47, 39–62.

Langmead B, Trapnell C, Pop M, Salzberg SL. 2009. Ultrafast and memory-efficient alignment of short DNA sequences to the human genome. Genome Biology 10, R25.

Laufs P, Peaucelle A, Morin H, Traas J. 2004. MicroRNA regulation of the CUC genes is required for boundary size control in Arabidopsis meristems. Development 131, 4311–4322.

Lee MH, Jeon HS, Kim HG, Park OK. 2017. An Arabidopsis NAC transcription factor NAC4 promotes pathogen-induced cell death under negative regulation by microRNA164. New Phytologist 214, 343–360.

Li J, Guo G, Guo W, Guo G, Dan T, Ni Z, Sun Q, Yao Y. 2012. miRNA164-directed cleavage of ZmNAC1 confers lateral root development in maize (Zea mays L.). BMC Plant Biology 12, 220.

Li S, Liu L, Zhuang X, et al. 2013. MicroRNAs inhibit the translation of target mRNAs on the endoplasmic reticulum in Arabidopsis. Cell 153, 562–574.

Li Y, Lu YG, Shi Y, et al. 2014. Multiple rice microRNAs are involved in immunity against the blast fungus Magnaporthe oryzae. Plant Physiology 164, 1077–1092.

Liu H, Cheng Q, Zhe C, Tao Z, Yang X, Zhou H. 2014. Identification of miRNAs and their target genes in developing maize ears by combined small rna and degradome sequencing. BMC Genomics 15, 25.

Liu Y, Nakayama N, Schiff M, Litt A, Irish VF, Dineshkumar SP. 2004. Virus induced gene silencing of a deficiens ortholog in Nicotiana benthamiana. Plant Molecular Biology 54, 701–711.

Mallory AC, Dugas DV, Bartel DP, Bartel B. 2004. MicroRNA regulation of NAC-domain targets is required for proper formation and separation of adjacent embryonic, vegetative, and floral organs. Current Biology 14, 1035–1046.

Mao X, Cai T, Olyarchuk JG, Wei L. 2005. Automated genome annotation and pathway identification using the KEGG Orthology (KO) as a controlled vocabulary. Bioinformatics 21, 3787–3793.

Mi S, Cai T, Hu Y, et al. 2008. Sorting of small RNAs into Arabidopsis argonaute complexes is directed by the 5′ terminal nucleotide. Cell 133, 116–127.

Navarro L, Dunoyer P, Jay F, Arnold B, Dharmasiri N, Estelle M, Voinnet O, Jones JD. 2006. A plant miRNA contributes to antibacterial resistance by repressing auxin signaling. Science 312, 436–439.

Nuruzzaman M, Manimekalai R, Sharoni AM, Satoh K, Kondoh H, Ooka H, Kikuchi S. 2010. Genome-wide analysis of NAC transcription factor family in rice. Gene 465, 30–44.

Ooka H, Satoh K, Doi K, Nagata T, et al. 2003. Comprehensive analysis of NAC family genes in Oryza sativa and Arabidopsis thaliana. DNA Research 10, 239–247.

Pang J, Zhu Y, Li Q, Liu J, Tian Y, Liu Y, Wu J. 2013. Development of Agrobacterium-mediated virus-induced gene silencing and performance evaluation of four marker genes in Gossypium barbadense. PLoS One 8, e73211.

Rajagopalan R, Vaucheret H, Trejo J, Bartel DP. 2006. A diverse and evolutionarily fluid set of microRNAs in Arabidopsis thaliana. Genes & Development 20, 3407–3425.

Schwechheimer C, Zourelidou M, Bevan MW. 1998. Plant transcription factor studies. Annual Review of Plant Physiology and Plant Molecular Biology 49, 127–150.

Sha A, Zhao J, Yin K, Tang Y, Wang Y, Wei X, Hong Y, Liu Y. 2014. Virus-based microRNA silencing in plants. Plant Physiology 164, 36–47.

Shi M, Hu X, Wei Y, Hou X, Yuan X, Liu J, Liu Y. 2017. Genome-wide profiling of small rnas and degradome revealed conserved regulations of miRNAs on auxin-responsive genes during fruit enlargement in peaches. Plant Physiology and Biochemistry 18, 2599.

Shriram V, Kumar V, Devarumath RM, Khare TS, Wani SH. 2016. MicroRNAs as potential targets for abiotic stress tolerance in plants. Frontiers in Plant Science 7, 817.

Sieber P, Wellmer F, Gheyselinck J, Riechmann JL, Meyerowitz EM. 2007. Redundancy and specialization among plant microRNAs: role of the MIR164 family in developmental robustness. Development 134, 1051–1060.

Sink KC, Grey WE. 1999. A root-injection method to assess verticillium wilt resistance of peppermint (mentha × piperita L.) and its use in identifying resistant somaclones of cv. black mitcham. Euphytica 106, 223–230.

Song QX, Liu YF, Hu XY, Zhang WK, Ma B, Chen SY. 2011. Identification of miRNAs and their target genes in developing soybean seeds by deep sequencing. BMC Plant Biology 11, 5.

Tamura K, Peterson D, Peterson N, Stecher G, Nei M, Kumar S. 2011. MEGA5: molecular evolutionary genetics analysis using maximum likelihood, evolutionary distance, and maximum parsimony methods. Molecular Biology and Evolution 28, 2731–2739.

Varkonyi-Gasic E, Wu R, Wood M, Walton EF, Hellens RP. 2007. Protocol: a highly sensitive RT-PCR method for detection and quantification of microRNAs. Plant Methods 3, 12.

Wang C, He X, Wang X, Zhang S, Guo X. 2017a. ghr-miR5272a-mediated regulation of GhMKK6 gene transcription contributes to the immune response in cotton. Journal of Experimental Botany 68, 5895–5906.

Wang L, Wu SM, Zhu Y, Fan Q, Zhang ZN, Hu G, Peng QZ, Wu JH. 2017b. Functional characterization of a novel jasmonate ZIM-domain interactor (NINJA) from upland cotton (Gossypium hirsutum). Plant Physiology and Biochemistry 112, 152–160.

Wang Q, Liu N, Yang X, Tu L, Zhang X. 2016. Small RNA-mediated responses to low- and high-temperature stresses in cotton. Scientific Reports 6, 35558.

Wang YQ, Chen DJ, Wang DM, Huang QS, Yao ZP, Liu FJ. 2004. Over-expression of Gastrodia anti-fungal protein enhances Verticillium wilt resistance in coloured cotton. Plant Breeding 123, 454–459.

Wei LQ, Yan LF, Wang T. 2011. Deep sequencing on genome-wide scale reveals the unique composition and expression patterns of microRNAs in developing pollen of Oryza sativa. Genome Biology 12, R53.

Xie F, Wang Q, Sun R, Zhang B. 2015. Deep sequencing reveals important roles of microRNAs in response to drought and salinity stress in cotton. Journal of Experimental Botany 66, 789–804.

Xin M, Wang Y, Yao Y, Xie C, Peng H, Ni Z, Sun Q. 2010. Diverse set of microRNAs are responsive to powdery mildew infection and heat stress in wheat (Triticum aestivum L.). BMC Plant Biology 10, 123.

Yan J, Gu Y, Jia X, Kang W, Pan S, Tang X, Chen X, Tang G. 2012. Effective small RNA destruction by the expression of a short tandem target mimic in Arabidopsis. The Plant Cell 24, 415–427.

Yin Z, Li Y, Han X, Shen F. 2012. Genome-wide profiling of miRNAs and other small non-coding RNAs in the Verticillium dahliae-inoculated cotton roots. PLoS One 7, e35765.

Yip HK, Floyd SK, Sakakibara K, Bowman JL. 2016. Class III HD-Zip activity coordinates leaf development in Physcomitrella patens. Developmental Biology 419, 184–197.

Yu F, Huaxia Y, Lu W, Wu C, Cao X, Guo X. 2012. GhWRKY15, a member of the WRKY transcription factor family identified from cotton (Gossypium hirsutum L.), is involved in disease resistance and plant development. BMC Plant Biology 12, 133.

Zhang T, Zhao YL, Zhao JH, Wang S, Jin Y, Chen ZQ, Fang YY, Hua CL, Ding SW, Guo HS. 2016. Cotton plants export microRNAs to inhibit virulence gene expression in a fungal pathogen. Nature Plants 2, 16153.

Zhang Y, Wang W, Chen J, Liu J, Xia M, Shen F. 2015a. Identification of miRNAs and their targets in cotton inoculated with verticillium dahliae by high-throughput sequencing and degradome analysis. International Journal of Molecular Sciences 16, 14749–14768.

Zhang Z, Jiang L, Wang J, Gu P, Chen M. 2015b. MTide: an integrated tool for the identification of miRNA-target interaction in plants. Bioinformatics 31, 290–291.

Zhao JP, Jiang XL, Zhang BY, Su XH. 2012. Involvement of microRNA-mediated gene expression regulation in the pathological development of stem canker disease in Populus trichocarpa. PLoS One 7, e44968.

Zhu QH, Fan L, Liu Y, Xu H, Llewellyn D, Wilson I. 2013. miR482 regulation of NBS-LRR defense genes during fungal pathogen infection in cotton. PLoS One 8, e84390.

